# 2B4 co-engagement promotes serial degranulation and killing by human NK cells

**DOI:** 10.64898/2026.07.31.741999

**Authors:** Luca Kröll, Mina Sandusky, Michèle Saretzki, Maren Claus, Sabine Wingert, Jens Niemann, Carsten Watzl

## Abstract

Activation of Natural Killer (NK) cells depends on the stimulation of a broad range of receptors. Here we use murine NIH3T3 target cells expressing defined human NK cell ligands to stimulate different aspects of NK cell activity and identify how distinct ligand-receptor interactions regulate different aspects of NK cell activation. B7H6, MICA, and CD20-bound obinutuzumab engaging NKp30, NKG2D, or CD16, respectively, were found to be potent inducers of activity in pre-stimulated NK cells. Combination of these ligands with CD48 to also stimulate 2B4 significantly increased degranulation and cytokine secretion, while PVR engaging DNAM-1 showed favorable effects only in combination with B7H6. Activating NK cell receptors were downregulated in a ligand-specific fashion, whereas CD16 and NKp46 were found to be downregulated depending on NK cell activation induced by other receptors. Co-stimulation *via* 2B4 in combination with either NKp30, NKG2D, or CD16 significantly enhanced NK cells to serially degranulate and kill multiple targets. Our systematic analysis unravels the complexity of different activating NK cell receptors and provides strategies to specifically tune NK cell reactivities for therapeutic applications.

## Introduction

Natural Killer (NK) cells are innate lymphocytes with an important role in early immune responses against virally infected or transformed cells (Cerwenka and Lanier, 2001). As effector cells, their main functions are the secretion of cytokines and chemokines to coordinate other players of the immune system and cytotoxic activity to eliminate compromised cells (Cooper et al., 2001). While other lymphocytes require prior sensitization with highly specific antigens, NK cell activation depends on the balance of activating and inhibitory signals from a plethora of germline-encoded receptors (Vivier et al., 2004, Watzl and Long, 2010). Based on their structural properties, associated signaling proteins, and functional outcome, activating receptors can be classified into different groups.

NKp30 (CD337), NKp44 (CD336), and NKp46 (CD335) are members of the natural cytotoxicity receptors (NCRs) (Moretta et al., 2000). While a cellular ligand for NKp46 remains unknown to date, B7H6 has been described as NKp30-ligand that is not abundantly expressed in normal tissue but frequently upregulated in different forms of cancer (Brandt et al., 2009). Upon engagement by its ligand and subsequent receptor crosslinking, NKp30 associates with adapter proteins containing immunoreceptor tyrosine-based activation motifs (ITAMs) to propagate activating signals within the cell (Watzl and Urlaub, 2012). In a similar manner, signaling of low-affinity IgG receptor CD16 (FcγRIII) depends on ITAM-containing adapters (Leibson, 1997, Watzl and Urlaub, 2012). Due to its recognition of the Fc region of antibodies, CD16 enables NK cells to mediate antibody-dependent cellular cytotoxicity (ADCC). The ability to engage with and kill opsonized cells plays a major role in antibody-based therapies against cancers or in autoimmune diseases (Natsume et al., 2009). One example for this includes CD20-directed obinutuzumab (obi) which was designed to overcome resistances that developed during treatment of non-Hodgkin lymphomas with rituximab (Freeman and Sehn, 2018, Rezvani and Maloney, 2011). NKG2D (CD314) is an NK cell activating receptor that can bind to MICA, MICB, and members of the ULBP protein family (Lanier, 2015). Similar to B7H6, these proteins are found to be significantly upregulated in various cancers (Raulet et al., 2013). To transduce activating signals upon binding, NKG2D forms a complex with adapter protein DAP10 (Garrity et al., 2005). DAP10 employs an immunoglobulin tyrosine tail (ITT) motif (YINM) to confer downstream signaling (Watzl and Urlaub, 2012).

Receptors belonging to the signaling lymphocytic activation molecule (SLAM)-family feature an immunoreceptor tyrosine-based switch motif (ITSM) in their cytosolic domain (Claus et al., 2008). This allows members of this group of receptors, which includes 2B4 (CD244), NTB-A, and CRACC, among others, to propagate signals independently. CD48 is the cellular ligand of 2B4. It is found on the surface of various hematopoietic cells and has been shown to be upregulated in viral infections and upon interferon (IFN) stimulation (Tissot et al., 1997, Klaman and Thorley-Lawson, 1995). DNAM-1 (CD226) is described as another (co-)activating receptor on NK cells (Tahara-Hanaoka et al., 2004, Bottino et al., 2003). Its ligands PVR (CD155) and Nectin-2 (CD112) are expressed on stressed, virus-infected, or transformed cells (Kamran et al., 2013, Li et al., 2018). DNAM-1, however, is not the only receptor expressed on NK cells found to bind to these ligands. With different affinities, inhibitory receptors TIGIT and CD96 compete with DNAM-1 over ligand binding (Samanta and Almo, 2015). The relationship and balance of these receptors have functional implications i.e. in hepatocellular carcinoma (Sun et al., 2019). In order to transduce activating signals, DNAM-1 has been shown to be dependent on the association with adhesion molecule LFA-1 (CD11a-CD18) (Shibuya et al., 1999). Upon contact with its ligand ICAM-1 on cells, LFA-1 switches its confirmation. This interaction has shown not only to provide strong adhesion but also to prime NK cell activity by mediating granule polarization (Barber et al., 2004).

When NK cells recognize viable target cells through the up-regulation of activating ligands (“induced-self”) or the lack of inhibitory ligands (“missing-self”) (Moretta et al., 2004), target cell death can be induced by two distinct pathways (Prager and Watzl, 2019). On the one hand, NK cells can secrete cytotoxic granules containing granzymes and perforin as effector molecules. These granules further contain CD107a (LAMP-1), whose presence on the surface of NK cells can be used to specifically detect degranulating cells. On the other hand, NK cells up-regulate death ligands TRAIL and Fas ligand (FasL) on their surface. Engagement of the respective receptors TRAIL receptor (TRAILR) or Fas on target cells induces a caspase-dependent signaling cascade resulting in apoptosis. During serial killing, NK cells have been shown to switch from granzyme-mediated to death receptor-mediated cytotoxicity (Prager et al., 2019). Additionally, NK cells can be triggered to secrete various cytokines and chemokines (Cooper et al., 2001, Bryceson et al., 2006a). This includes, among others, IFN-γ, TNF, and GM-CSF. Secretion of these cytokines helps coordinating immune responses especially during early phases of infection.

The diversity of activating receptor families, their individual signal transduction pathways, and the involvement of inhibitory signals renders NK cell stimulation highly complex. Several synergies and relationships between receptors have been observed under different conditions. This includes, for example, the synergy between NKG2D, 2B4, and LFA-1 that was subsequently attributed to NFκB signaling (Bryceson et al., 2009, Kwon et al., 2016). Other examples include the cooperation between LFA-1 and DNAM-1 (Shibuya et al., 1999) or the synergy between NKp46 and 2B4 (Zamai et al., 2020). However, most studies focusing on the detailed nature of NK cell activation processes were conducted using *drosophila* S2 cells expressing defined ligands for human NK cell receptors, plate-bound antibodies, or coated microbeads as targets. These *in vitro* model systems strongly deviate from physiological conditions and therefore fail to faithfully recapitulate the complex interactions of NK and tumor cells. In this work, we use murine embryonal NIH3T3 (3T3) cells as mammalian cellular targets enabling prolonged incubation times under optimal conditions for both target and effector cells. By expressing the human NK cell ligands B7H6, CD48, PVR, MICA, Nectin-2, ICAM-1, or CD20, different target cell lines were generated. Because wild type (wt) 3T3 cells do not specifically interact with human NK cells, activity observed with transfected cells could be pinned down to the interaction of specific ligand-receptor pairings, allowing for the systematic identification of distinct cause-effect relationships. Our data show that NKp30, NKG2D and CD16 are best at stimulating NK cell responses that can only be slightly enhanced by ICAM-1 co-expression. While PVR can enhance B7H6-mediated responses, 2B4 co-engagement can not only enhance NKG2D-, CD16-, and NKp30-mediated NK cell activation, but specifically boost serial degranulation and killing.

## Methods

### NK cell isolation

NK cells were isolated from peripheral blood mononuclear cells (PBMCs) of healthy donors by negative selection using the Dynabeads™ Untouched™ Human NK cells kit (Invitrogen™, Thermo Fisher Scientific) according to the manufacturer’s protocol. All blood donors gave informed consent and the study was approved by the Ethics Committee of the Leibniz Research Center (#213). Purity of NK cells was verified by staining for CD3 and CD56 using flow cytometry.

### NK cell culture

Freshly isolated NK cells were co-cultured with irradiated K562-41BBL-mbIL-15-mbIL-21 feeder cells at a ratio of 2:1 (NK:K562) in a 96-well round-bottom-plate in IMDM GlutaMax™ (Gibco®, Thermo Fisher Scientific) with 10 % (v/v) Fetal Bovine Serum (FBS; Gibco®, Thermo Fisher Scientific), 1 % (v/v) Penicillin/Streptomycin (P/S; Gibco®, Thermo Fisher Scientific), and 200 U/mL IL-2 (NIH Cytokine Repository). Medium was regularly replaced and supplemented with 100 U/mL IL-2. When NK cells exceeded 3 · 10^6^ cells/mL they were diluted to 1.5 · 10^6^ cells/mL. Assays in this work were performed with NK cells that had been cultured for at least 21 and at most 42 days.

### NIH3T3 cell culture and transfection

NIH3T3 cell lines were cultured in DMEM (Gibco®, Thermo Fisher Scientific) with 10 % FBS and 1 % P/S (3T3 medium). Cells were transfected using Lipofectamine™ 2000 (Invitrogen™, Thermo Fisher Scientific) and selected using 1 µg/mL Puromycin and/or 1 mg/mL G418. Purity of transfected cell lines was confirmed by staining for respective surface proteins using flow cytometry and cells were further enriched by cell sorting (BD FACSAria™ Fusion). When using CD20-expressing NIH3T3 cells for functional experiments, cells were incubated with 1 µg/mL obinutuzumab (obi) before the assay.

### HeLa cell culture

HeLa cells and HeLa cells stably transfected with CD48 (HeLa-CD48) were maintained in 3T3 medium supplemented with 0.5 µg/mL puromycin in case of CD48 transfection.

### Conjugate formation assay

NIH3T3 and NK cells were washed with serum free 3T3 medium and labeled for 30 min with 0.2 µM CellTracker™ DeepRed and 1 µM CellTrace™ CFSE (both Thermo Fisher Scientific), respectively. Cells were then washed, resuspended in 3T3 medium without phenol red, transferred to BIOFLOAT™ 96-well round-bottom plates (Sarstedt) at an effector-to-target (E/T) ratio of 2:1, centrifugated for 30 s at 100 x *g,* and incubated at 37 °C. To stop reactions at the desired time points, samples were fixed with an equal volume of ice-cold 4 % polyformaldehyde solution and repeatedly resuspended. Conjugates were analyzed by flow cytometry on a BD LSRFortessa® and data were analyzed using FlowJo™ Software (BD Life Sciences).

### Degranulation assay with NIH3T3 targets

NK cells were labeled with 0.2 µM CellTracker™ DeepRed, washed, and incubated for 20 min at 37 °C in IMDM with 10 % FBS and 1 % P/S (CTL medium). Cells were washed again, transferred to a 96-well V-bottom plate at an E/T of 1:1 and incubated in the presence of a BV421-conjugated anti-CD107a antibody for 3 h at 37 °C. After that, cells were washed and resuspended in DPBS + 2 % FBS (FACS buffer) for measurement. NK cell samples without target cells were used as control for spontaneous degranulation.

### Cytokine release assay

NIH3T3 cells were seeded in CTL medium in a 96-well flat-bottom plate (50,000 cells per well). After 6 hours, NK cells were added at an E/T ratio of 1:1 and the plate was incubated overnight. Plates were centrifugated and supernatants were either collected for further processing or frozen at –80 °C for later measurements. Cytokines were measured in duplicates using a LEGENDPlex™ Human CD8/NK Panel (BioLegend). NK cell samples without NIH3T3 targets were employed to control for cytokine base levels.

### NK receptor panel

NIH3T3 and NK cells were co-incubated at a 1:1 ratio in a 96-well round-bottom plate for 3 hours in the presence of a PerCP-conjugated anti-CD107a antibody. Cells were washed, stained with Zombie yellow, washed again, and stained using a mix of antibodies against the receptors of interest. Flow cytometric measurements of these samples were performed with a Cytek® Aurora using NK cells without targets as controls.

### Serial degranulation

To detect NK cells degranulating against targets in a serial fashion we used a slightly modified assay which we recently developed (Niemann et al., 2025). In short, NIH3T3 and NK cells were co-incubated at an E/T of 1:4 for 15 min at 37 °C. Next, a fluorophore-coupled anti-CD107a antibody was added and cells were incubated for another 10 min. Finally, residual binding sites were blocked by adding an unlabeled anti-CD107a antibody in excess for 5 min before washing cells with CTL medium. This process was repeated twice with different fluorophore-coupled anti-CD107a antibodies. Serial degranulating NK cells were analyzed by flow cytometry as described (Niemann et al., 2025).

### 51Cr-release cytotoxicity assay

NIH3T3 cells were labeled for 1 h with Na2^51^CrO4 (revvity), washed and co-incubated with NK cells at different E/T ratios for 4 h at 37 °C in 3T3 medium. Incubation in medium without NK cells or with 2 % (v/v) Triton X-100 served as controls for spontaneous and maximal ^51^Cr-release from target cells, respectively. Plates were centrifugated and supernatants were transferred to tubes containing Perleen® ICE absorbent powder. Samples were measured in triplicates and radiation was read out with a Wizard^2^ gamma counter (Perkin Elmer). Specific lysis of target cells was determined as follows:

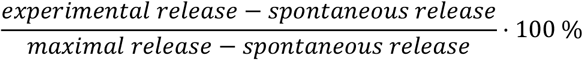

### Impedance-based cytotoxicity assay

16-well PET E-plate slides (OLS) were coated with 1.25 µg collagen (Sigma Aldrich) per well for 30 minutes. Plates were washed and 50 µL CTL medium was added to each well. After a short incubation at 37 °C, basic impedance levels were determined using an xCelligence RTCA DP analyzer (Agilent). Then, 50 µL of NIH3T3 cells at 1.6 · 10^6^ cells/mL were transferred to coated plates in duplicates. Impedance was tracked every 15 min for the next 24 h. After that, NK cells were added to target cells at an E/T ratio of 2:1. Measurement continued in 2 min intervals for the next 3 h, where NIH3T3 cell samples without NK cells served as control.

### Time lapse microscopy of NK cell serial killing

Microscopy was performed on a Mica WideFocal Live Cell microscope (Leica) with the inbuild 10x objective. HeLa and HeLa-CD48 cells were seeded into an 8-well microscopy dish at 2 · 10^5^ cells/well in 200 µL and left to adhere overnight at 37 °C. The next day, medium was exchanged with IMDM without phenol red containing 10 % FCS and 1 % P/S supplemented with 100 U/mL IL-2. For imaging, areas with uniform cell distribution were chosen and 2.5 · 10^3^ NK cells stained with CellTracker™ Green (Thermo Fisher Scientific) were added to each well. Images of brightfield and CellTracker™ Green channel were taken every 3 min for 16 h. Videos were analyzed manually.

### List of antibodies

PE anti-CD3 (BioLegend, #300308), BV421 anti-CD56 (BD BioSciences, #562752), PE anti-CD54 (BD BioSciences, #555511), PacificBlue anti-CD54 (BioLegend, #353109), PE anti-B7H6 (R&D Systems, #FAB7144P), APC anti-B7H6 (R&D Systems, #FAB7144A), PE anti-CD48 (BioLegend, #336707), PE anti-CD155 (BioLegend, #337609), PE anti-MICA/B (BioLegend, #320906), APC anti-MICA/B (BioLegend, #320907), APC anti-CD112 (BioLegend, #337411), PE anti-CD20 (BioLegend, #302305), APC anti-CD20 (BioLegend, #302309), PE anti-NKp30 (BioLegend, #325208), PE anti-NKG2D (BioLegend, #320806), PE anti-CD16 (BioLegend, #302008), PE anti-2B4 (BioLegend, #329508), AF647 anti-DNAM-1 (BioLegend, #338327), PE anti-TIGIT (BioLegend, #372703), PE anti-CD96 (BioLegend, #338405), AF488 anti-CD11a (BioLegend, #301216), AF700 anti-CD18 (BioLegend, #302124), BV421 anti-CD107a (BioLegend, #328625), BUV805 anti-CD56 (BD BioSciences, #742022), APC/Fire™750 anti-NKp30 (BioLegend, #325225), BV421 anti-NKp46 (BioLegend, #331913), PE-Dazzle594 anti-CD16 (BioLegend, #302054), PE-Cy5.5 anti-2B4 (Beckman Coulter, #B21171), BV786 anti-TIGIT (BD BioSciences, #747838, #570380), PerCP anti-CD107a (BioLegend, #328642), FITC anti-CD107a (BioLegend, #328605), SparkRed718 anti-CD107a (BioLegend, #328660).

## Results

### NKp30, CD16, and NKG2D show superior NK cell activating properties

To investigate specific receptor-ligand interactions between NK cells and targets, NIH3T3 cell lines expressing individual human NK cell ligands were generated (Fig. S1). We expressed B7H6 to engage NKp30, CD48 to stimulate 2B4, PVR or Nectin-2 as ligands for DNAM-1, MICA to stimulate NKG2D, ICAM-1 to engage LFA-1 and CD20 to use the anti-CD20 antibody obinutuzumab for the activation of CD16. Comparing expression levels of the newly introduced surface proteins with commonly employed human cancer cell lines showed that expression levels were within reasonable range for all generated NIH3T3 cells (Fig. S2). Additionally, variability of NK cell receptor expression levels was assessed by staining cultured human NK cells from different donors, revealing only minor donor-to-donor variations (Fig. S3). Following this initial establishment, NK cell ligand-expressing NIH3T3 cell lines were used to investigate how individual ligands stimulate cultured human NK cells using various readouts. Investigating conjugate formation, we found that target cells expressing adhesion ligand ICAM-1 rapidly and sustainably engage NK cells, forming more than three times as many conjugates after 10 min of co-incubation as NIH3T3 wt controls (Fig. 1A, B). ICAM-1 serves as ligand for adhesion receptor LFA-1 and was therefore expected to account for the largest fraction of conjugates. To a smaller extend, targets expressing B7H6, PVR, MICA, or CD20 showed significant but unsustainable and transient conjugate formation (Fig. 1A, B). In a flow cytometry-based degranulation assay, significant degranulation was induced after NK cells were co-incubated with any transfected target cell line (Fig. 1C). Interestingly, two distinct groups could be observed, where cells with NKp30-, NKG2D-, and CD16-engaging ligands emerged as superior inducers of degranulation as opposed to 2B4-, DNAM-1-, and LFA-1-targeting ligands. In ^51^Cr-release assays, however, only B7H6-expressing target cells could confirm these observations, clearly inducing NK cell cytotoxicity to the highest extend (Fig. 1D). Beyond that, incubation with remaining NIH3T3 cell lines led to mostly significant but less pronounced cytotoxic activity of different degrees. To validate these cytotoxicity results using a different experimental read-out, an impedance-based assay was performed. Here, B7H6-, MICA-, and CD20-expressing targets emerged as the only cells that significantly increased cytotoxicity within 3 hours of co-culture with NK cells (Fig. 1E). Interestingly, results for ICAM-1-expressing cells in this assay suggested a significant reduction in NK cell cytotoxicity. This could, however, likely be attributable to the adhesive properties of ICAM-1, which might have interfered with target cells properly detaching from the experimental vessel, thereby maintaining increased impedance. Because of this uncertainty, subsequent evaluations of cytotoxic activity were performed solely by ^51^Cr-release measurements. Finally, secretion of cytotoxic effectors and cytokines was evaluated using a bead-based flow cytometric approach. While no statistically significant increases were found for any target cell line, NK cells co-incubated with B7H6-expressing targets showed the most pronounced enhancement in IFN-γ and GrzB levels, followed by MICA-, Nectin-2-, and CD20-expressing cells (Fig. 1F).

**Figure 1:**
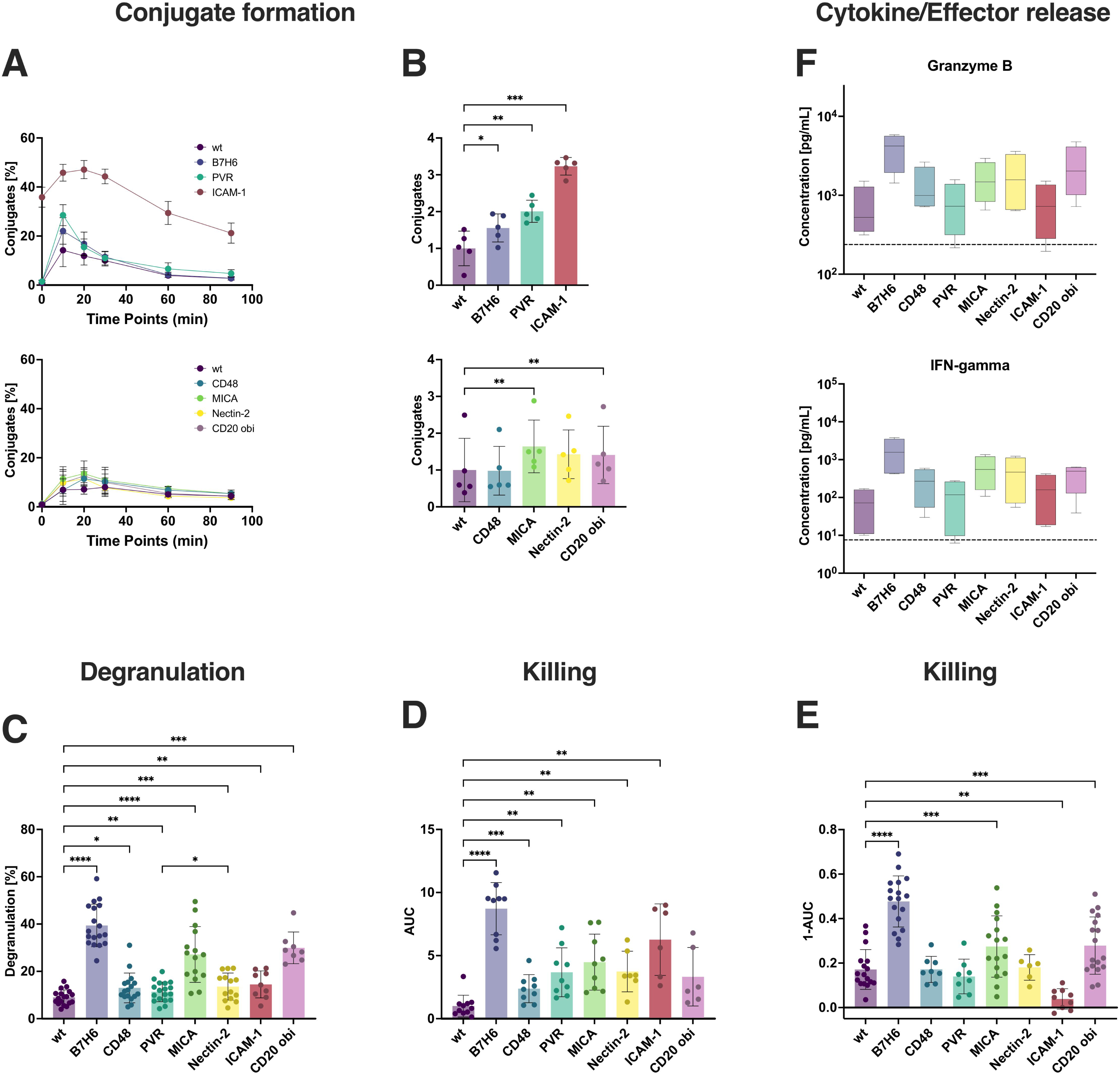
B7H6, MICA, and CD20 in combination with obinutuzumab are potent inducers of NK cell activity. NIH3T3 cells were transfected to express one of seven specific human NK cell ligands and subsequently co-incubated with pre-stimulated NK cells to investigate different aspects of NK cell activation. **A** Time-dependent formation of conjugates with respective target cell lines at an effector to target (E/T) ratio of 1:1; n = 5. obi: obinutuzumab. **B** Statistical comparison of conjugates formed with different targets after 10 minutes normalized to wt control. **C** Frequencies of CD107a^+^ NK cells after 3 hours of co-incubation at an E/T ratio of 1:1; n = 7-18. **D** Quantification of ^51^Cr-release cytotoxicity assays shown as AUC of fractions of dead target cells normalized to wt controls at E/T ratios of 8:1, 4:1, 2:1 and 1:1; n = 6-11. **E** Cytotoxicity assessed by impedance-based xCelligence assay. Results show AUC of cell index within 3 hours after co-incubation at an E/T ratio of 2:1 normalized to target cell controls without effectors; n = 6-17. **F** Secretion levels of GrzB and IFN-γ after overnight co-incubation at an E/T ratio of 1:1 with dashed lines indicating mean of NK cell controls without targets; n = 4. *Data information*: One-Way ANOVA with Dunnet (B, D, E, F) or Šidák correction (C). \**P* ≤ 0.05, \*\**P* ≤ 0.01, \*\*\**P* ≤ 0.001, \*\*\*\**P* ≤ 0.0001. Data presented as mean ± SD (A-E).

These initial results support NKp30-, NKG2D-, and CD16-engaging ligands as superior inducers of NK cell activities.

### Combination of activating ligands with ICAM-1 only leads to minor increases in NK cell functionality

The initial step in NK cell cytotoxicity is conducted by establishing and maintaining contact with its cellular target and forming an immunological synapse. In this process, adhesion receptor LFA-1 and its counterpart ICAM-1 are known to be centrally involved, which is reflected in the extensive conjugate formation capabilities of ICAM-1 expressing cells (Fig. 1A). To investigate the role of ICAM-1-mediated adhesion for NK cell activity, NIH3T3 cells expressing individual NK cell ligands were additionally transfected to co-express ICAM-1. Protein surface levels in co-expressing cell lines were comparable to those in cells expressing only one of the respective ligands (Fig. S1). Following the previously employed approach, NK cells were investigated regarding their capability to form stable conjugates with the newly generated target cell lines. While cells engaging only NKp30, NKG2D, or CD16 showed little increased conjugate formation when compared to wt controls, co-expression of ICAM-1 significantly elevated conjugation to levels comparable with ICAM-1 only expressing cells (Fig. 2A, B). This enhanced contact formation impacted subsequent degranulation and cytotoxicity assays to different degrees. When combined with NKp30-ligand B7H6, both degranulation and cytotoxic activity of NK cells were increased visibly, though not significantly, when compared to B7H6 only expressing controls (Fig. 2C, D). Co-engagement of NKG2D and LFA-1 led to no change in degranulation levels, but significantly increased NK cell activity in ^51^Cr-release assays. The largest impact of ICAM-1 co-expression, however, was observed for CD16. Here, frequencies of degranulating NK cells were almost doubled compared to controls without expression of ICAM-1 (Fig. 2C). While cytotoxicity was visibly increased against co-expressing targets as well, no statistical significance was observed in this case, which is likely attributed to the high scattering of obtained data (Fig. 2D). Combination of ICAM-1 with DNAM-1 ligands PVR or Nectin-2 or with the 2B4 ligand CD48, respectively, led to no remarkable changes in either degranulation or cellular cytotoxicity (Fig. S4).

**Figure 2:**
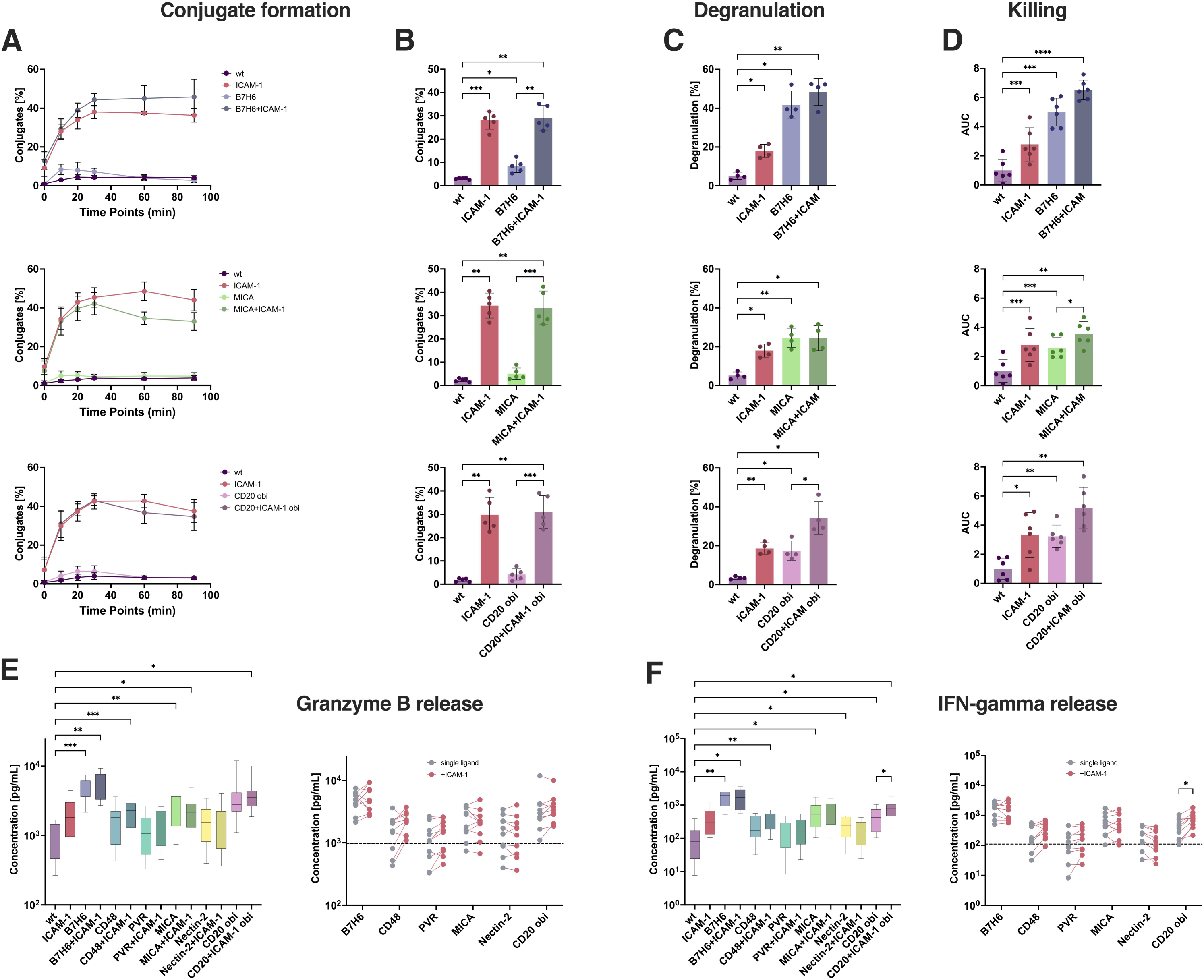
Adhesion ligand ICAM-1 in combination with other ligands has minor influence on NK cell stimulation. Assessment of NK cell activity after co-incubation with target cells co-expressing ICAM-1 and one of six other ligands. **A** Time-dependent formation of conjugates at an E/T ratio of 1:1; n = 5. **B** Fractions of NK cells conjugated with target cells after 10 min. **C** Fractions of CD107a^+^ NK cells after 3 hours of target cell co-incubation at an E/T ratio of 1:1; n = 4. **D** Quantification of NK cell cytotoxicity shown as AUC of dead target cells [%] at E/T ratios of 8:1, 4:1, 2:1, 1:1 normalized to wt controls; n = 6. **E** GrzB and **F** IFN-γ secretion levels after overnight co-incubation at an E/T ratio of 1:1, dashed lines indicate mean of wt controls; n = 10. *Data information*: One-Way ANOVA with Šidák correction (B, C, D, E left panel, F left panel) or Two-Way ANOVA with Šidák correction (E right panel, F right panel). \**P* ≤ 0.05, \*\**P* ≤ 0.01, \*\*\**P* ≤ 0.001, \*\*\*\**P* ≤ 0.0001. Data presented as mean ± SD (A-D).

Cooperation between CD16 and LFA-1 has previously been described (Bryceson et al., 2005) and was further highlighted when NK cell cytokine and effector molecule secretion were examined following overnight co-incubation with respective target cells (Fig. 2E, F). While GrzB secretion was not significantly increased for any cells co-expressing ICAM-1 compared to single ligand controls, IFN-γ was significantly enhanced against CD16-targeting cells.

Together, these results suggest that by LFA-1 co-engagement, NKp30– or NKG2D-mediated NK cell responses can be slightly increased, whereas CD16-induced responses are significantly enhanced by LFA-1 signals.

### Co-engagement of 2B4 *via* CD48 strongly enhances NK cell activation

2B4 and DNAM-1 are described as two major co-stimulating receptors on NK cells that can enhance NK cell activities in combination with other activating receptors but not trigger significant activation on their own (Long et al., 2013, Sivori et al., 2000). Therefore, we generated NIH3T3 cell lines that co-expressed either 2B4-ligand CD48 or DNAM-1-ligand PVR together with activating ligands B7H6, MICA, or CD20, respectively. Expression levels of individual proteins of newly generated cells were largely comparable to those of previously employed targets expressing only one of the respective ligands (Fig. S5). When CD48 was additionally expressed on target cells, degranulation against cells co-expressing MICA or CD20, respectively, was strongly increased in a synergistic fashion, while no further enhancement could be observed for B7H6-expressing targets (Fig. 3A). However, the 2B4-mediated enhancement was not observed in ^51^Cr-release cytotoxicity assays. While cytotoxic activity was slightly but significantly increased against MICA-co-expressing targets, no relevant changes were observed for NK cells co-incubated with B7H6– or CD20-expressing cells, respectively (Fig. 3B).

**Figure 3:**
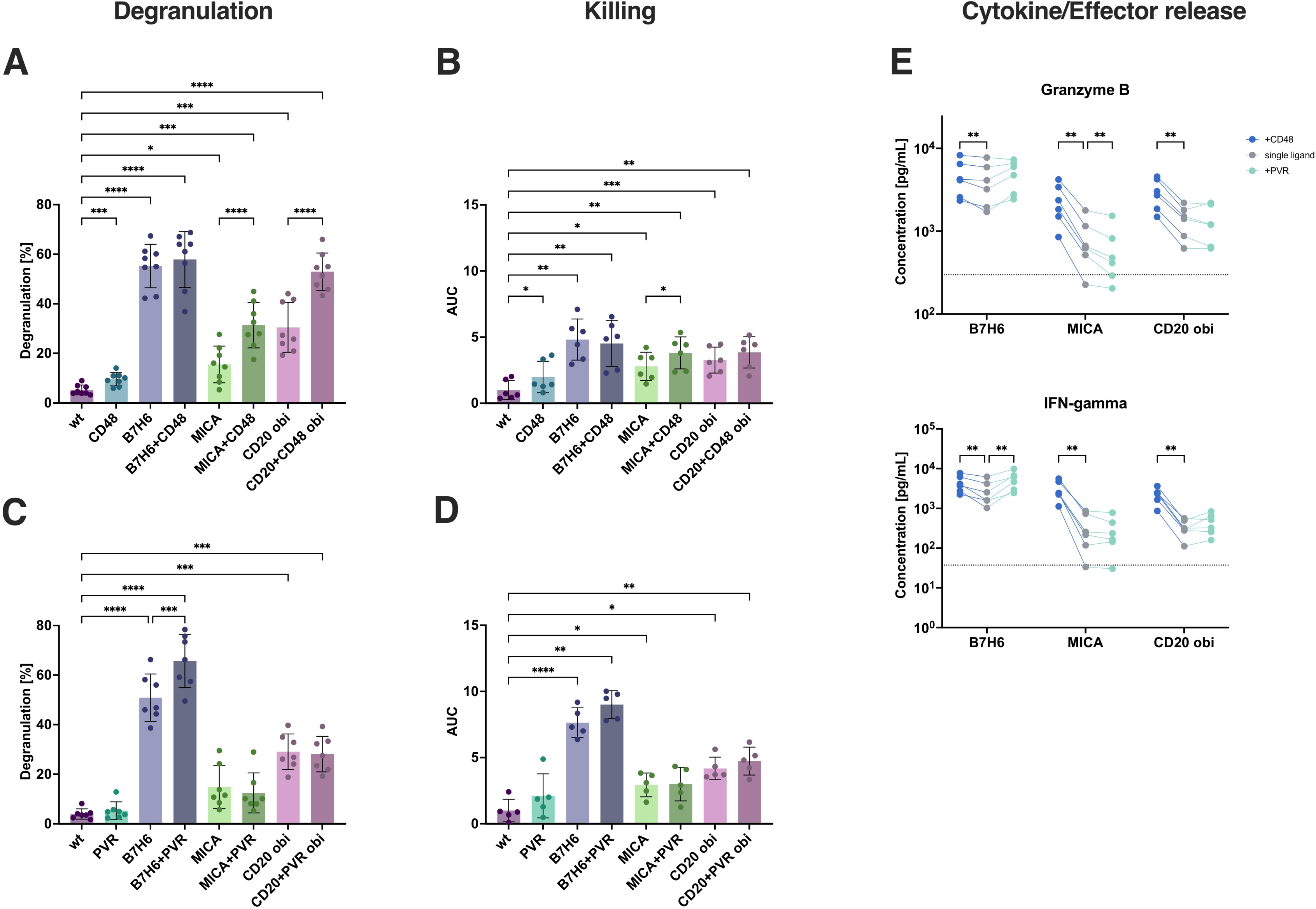
Co-engagement of 2B4 or DNAM-1 in combination with different other receptors strongly increases NK cell degranulation and effector and cytokine secretion. Comparison of NK cell stimulation by target cells expressing either single activating ligands or co-expressing co-stimulating ligands in addition. **A** Frequencies of degranulating NK cells after 3 hours of co-incubation with targets co-expressing CD48 at an E/T ratio of 1:1; n = 8. **B** NK cell cytotoxicity depicted as AUC of dead target cells [%] at E/T ratios 8:1, 4:1, 2:1, 1:1 in ^51^Cr-release assays; n = 6. **C-D** Same as A-B for target cells co-expressing PVR instead of CD48; n = 7 (C) and n = 5 (D). **E** Secretion levels of GrzB (top) and IFN-γ (bottom) after overnight incubation at an E/T ratio of 1:1 with dashed lines indicating means of wt controls; n = 6. *Data information*: One-Way ANOVA with Šidák correction (A-D) or Two-Way ANOVA with Dunnet correction (E). \**P* ≤ 0.05, \*\**P* ≤ 0.01, \*\*\**P* ≤ 0.001, \*\*\*\**P* ≤ 0.0001. Data presented as mean ± SD (A-D).

When PVR was co-expressed on target cells a different picture emerged. Here, degranulation was significantly increased against B7H6-co-expressing targets, while no changes were observed for MICA– or CD20-co-expressing cells (Fig. 3C). Similarly, a small increase in cytotoxic activity was only observed after co-engagement of NKp30 and DNAM-1 (Fig. 3D).

Finally, target cells co-expressing co-stimulating ligands were investigated regarding their ability to induce secretion of effector molecules and cytokines by NK cells. Co-expression of PVR generally did not enhance the secretion induced by MICA or CD20 but slightly increased NKp30-mediated secretion of IFN-γ, TNF, and GrzA. In contrast, 2B4 co-engagement by CD48 caused NK cells to increase secretion of various effectors and cytokines, including GrzB, IFN-γ, and TNF, which was particularly pronounced in combination with MICA or CD20 (Fig. 3E, Fig. S6). The 2B4-mediated co-stimulation of B7H6-induced secretion was much less pronounced, possibly as B7H6-only induced secretion was already quite high.

These data show that PVR can enhance NKp30-mediated NK cell responses, whereas 2B4 seems to be best at enhancing NKG2D or CD16-induced activities.

### NK cell receptor levels can be reduced independent of activation

Upon extensive interaction with their physiological ligands, NK cell receptors can be downregulated due to internalization or shedding (Srpan et al., 2018, Ogasawara et al., 2003, Semeraro et al., 2015, Braun et al., 2020, Sandusky et al., 2006). Therefore, we determined relative surface levels of receptors after co-incubation with target cells and linked this to degranulation by flow cytometry (Fig. S7). Depending on ligand expression on target cells, NK receptor levels were affected differently and largely specifically. When NK cells were induced to degranulate against B7H6-expressing targets, NKp30 levels were markedly reduced (Fig. 4A). Similarly, DNAM-1, NKG2D, CD16, and 2B4 levels were decreased when exposed to PVR-, MICA-, CD20-, or CD48-expressing targets, respectively. Interestingly, Nectin-2-expressing targets did not cause DNAM-1 levels to drop in a similar way, nor did it result in a downregulation of TIGIT, in contrast to PVR. A recent study on the influence of DNAM-1-ligand levels on NK cell activity could confirm differential effects of PVR and Nectin-2 on DNAM-1 levels and NK cell activity (Saunders et al., 2026). Also, LFA-1 levels remained unchanged upon ICAM-1 expressing target cell contact. Interestingly, this ligand-specific downregulation of activating NK cell receptors could also be observed on non-degranulating NK cells (Fig. 4B), suggesting that degranulation and receptor downregulation are uncoupled, and contact with respective cellular ligands seems sufficient to cause receptor downregulation on NK cell surfaces for certain receptor-ligand pairs, regardless of subsequent activity. Using target cells co-expressing CD48 or PVR confirmed these results of ligand-specific receptor downregulation on degranulating but also on non-degranulating cells (Fig. 4C, D) and downregulation of 2B4 was more pronounced when CD48 was co-expressed together with either B7H6, MICA, or CD20.

**Figure 4:**
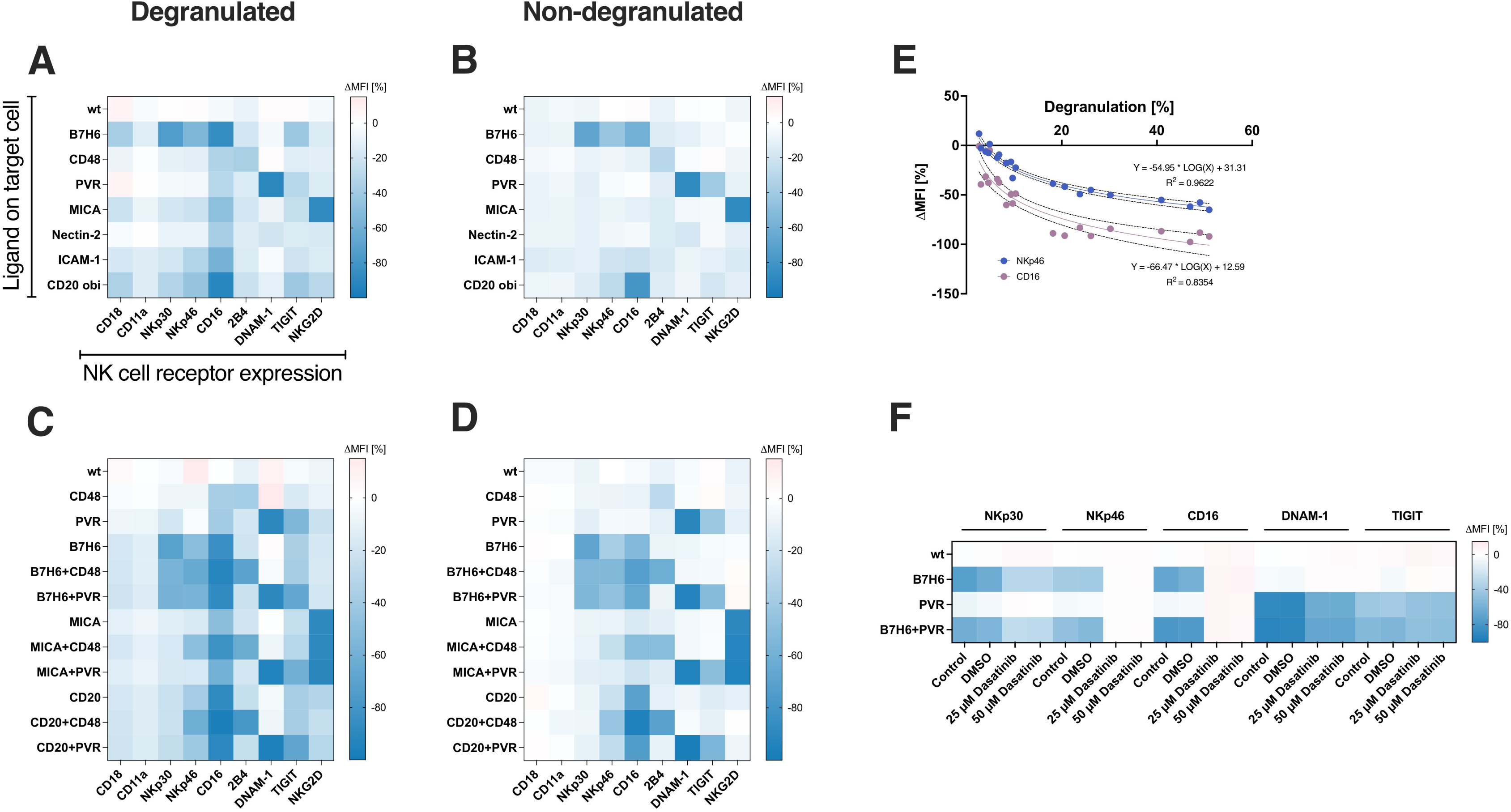
NK cell receptor levels are downregulated differently depending on ligand engagement, activating signals, and degranulation. Investigation of changes in expression levels of receptors on NK cell surfaces after co-incubation with different target cell lines compared to NK cell controls without target cells. **A** Percentual changes in MFI of fluorescently stained receptors after co-incubation for 3 hours with target cells expressing one of seven ligands on NK cells that either degranulated or **B** did not degranulate; n = 6. **C** Percentual changes of receptor expression levels after co-incubation with target cells co-expressing co-stimulating ligands CD48 or PVR on NK cells discriminated for degranulating and **D** non-degranulating cells; n = 6. **E** Correlation of frequencies of degranulating NK cells and changes in CD16 or NKp46 surface levels in these cells using data shown in panels A and C. **F** Percentual downregulations of NK cell receptors after treatment with tyrosine kinase inhibitor dasatinib and co-incubation with a selection of different target cell lines; n = 5. *Data information*: Semilogarithmic interpolation curve with 95 % confidence bands (E).

Some receptors were downregulated even when they were not engaged by ligands on the target cells. CD16 was downregulated by degranulating cells co-incubated with B7H6-, CD48-, PVR-, MICA-, Nectin-2, and ICAM-1-expressing target cells, confirming the previously described activation-induced shedding of the receptor (Srpan et al., 2018). Interestingly, NKp46 was also downregulated after stimulation via B7H6 or CD20, and CD48 co-expression could enhance this downregulation when combined with B7H6, CD20, and also MICA. When determining the amount of NKp46 and CD16 downregulation and the fraction of degranulating cells we found a clear correlation that could be fitted semi-logarithmically, with coefficients of determination of R^2^ = 0.9622 and R^2^ = 0.8354, respectively (Fig. 4E). This correlation was particularly interesting for NKp46, since there is no known ligand expressed on the NIH3T3 target cells.

Surprisingly, downmodulation of CD16 and NKp46 could also be observed on non-degranulating cells (Fig. 4B, D), which would contradict a purely activation-induced downregulation of these receptors. To directly investigate this, NK cells were treated with tyrosine kinase inhibitor dasatinib to block activation and subsequently co-incubated with different target cell lines. Downregulations of NKp46 and CD16 were completely abolished upon dasatinib treatment (Fig. 4H, Fig. S8). On the other hand, NKp30 and DNAM-1 levels were downregulated to a lesser extent upon treatment, while changes in TIGIT expression levels remained unchanged compared to DMSO and untreated controls. This indicates that engagement of NKp30, DNAM-1, or TIGIT by their specific cellular ligands alone causes NK cells to downregulate these receptors, while downregulation of NKp46 and CD16 is a by-product of activation by other receptors that does not occur when activating signals are blocked.

### Co-engagement of 2B4 promotes serial degranulation and killing

NK cells possess the ability to degranulate against and kill several target cells in a serial fashion (Bhat and Watzl, 2007). To investigate whether the fraction of serially degranulating NK cells depends on the engagement of specific receptors, NK cells were incubated with different target cell lines in a flow cytometry-based assay we previously established (Niemann et al., 2025). NK cells co-incubated with B7H6-expressing target cells showed the largest fraction of degranulating cells, followed by CD20– and MICA-expressing targets, respectively (Fig. 5A). Further, engagement of NKp30 caused the largest fraction of individual NK cells to degranulate against multiple targets (Fig 5B). Interestingly, CD16 levels on NK cells were reduced significantly depending on the number of degranulation events, independently of the stimulating receptor (Fig. 5C), confirming previous observations (Niemann et al., 2025). A similar trend was obtained for NKp46 surface levels (Fig. S9). However, results obtained for cells degranulating three times against MICA– or CD20-expressing need to be handled with caution, since this fraction made up less than 0.1 % of analyzed cells, which also explains the observed variability in data points.

**Figure 5:**
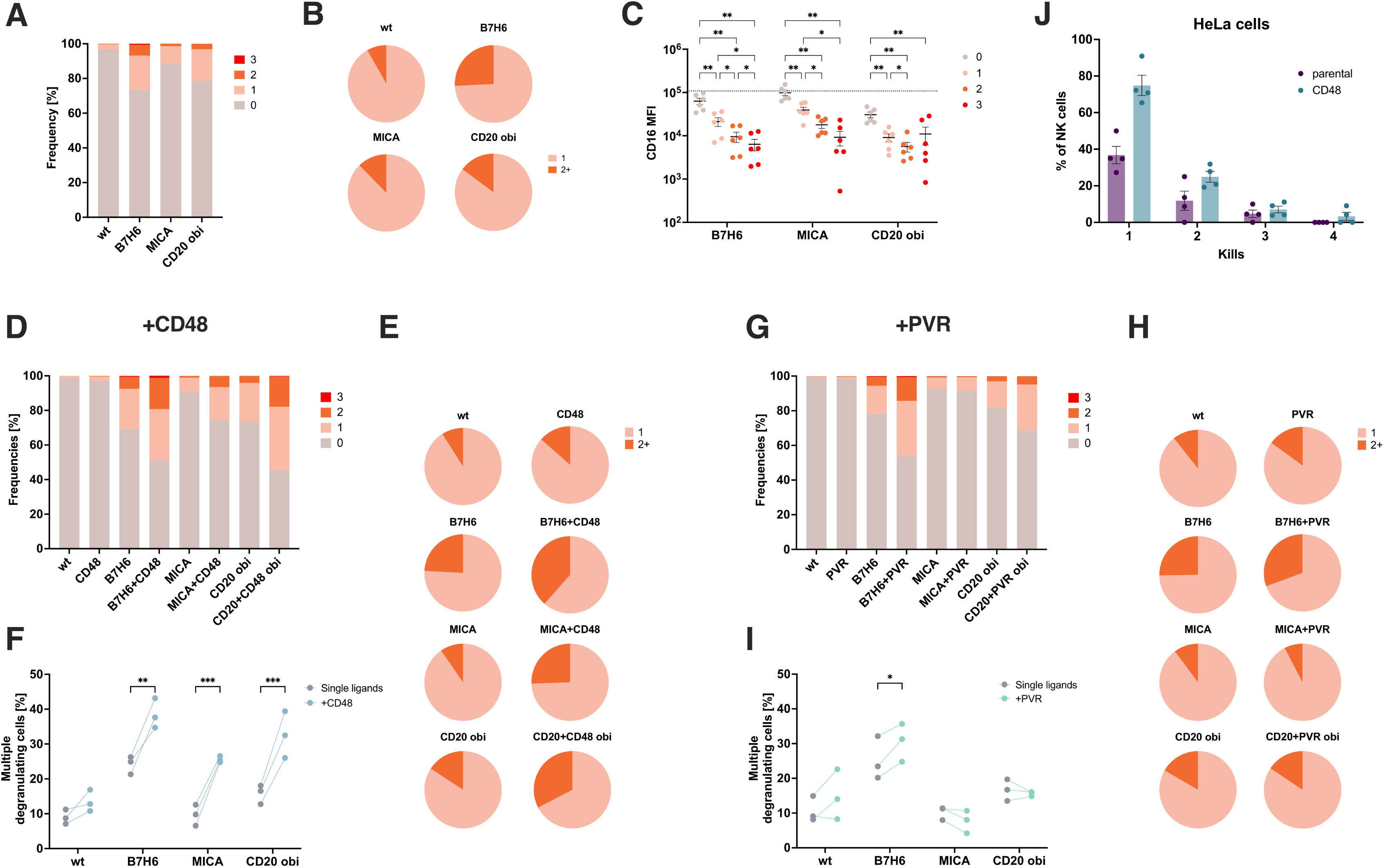
Serial degranulation in NK cells is enhanced significantly against target cells co-expressing 2B4-ligand CD48. Flow cytometry-based assessment of NK cells degranulating against multiple targets in serial fashion. **A** Frequencies of NK cells co-incubated with cellular targets expressing different ligands distributed by the number of degranulation events per cell; n = 8. **B** Pie charts showing the fraction of NK cells degranulating multiple times within the population of degranulating cells. **C** Receptor expression levels of CD16 on NK cells depending on the number of recorded degranulation events with dashed line indicating mean CD16 level in NK cell controls without targets; n = 6. **D** Frequencies of NK cells distributed by the number of degranulation events per cell after co-incubation with cellular targets co-expressing CD48. **E** Pie charts and **F** statistical quantification of the fractions of NK cells degranulating multiple times against CD48-co-expressing target cells; n = 3. **G-I** Same as D-F for target cells co-expressing PVR instead of CD48; n = 3. **J** Frequencies of NK cells grouped by the number of killing events against parental or CD48-expressing HeLa cells as determined by time lapse microscopy. *Data information*: Two-Way ANOVA with Tukey (C) or Šidák correction (F, I). \**P* ≤ 0.05, \*\**P* ≤ 0.01, \*\*\**P* ≤ 0.001, \*\*\*\**P* ≤ 0.0001. Data presented as mean ± SEM (C).

When analyzing targets co-expressing CD48, overall degranulation levels were increased visibly (Fig. 5D). Further, co-engagement of 2B4 caused fractions of serially degranulating cells to increase significantly across tested target cell lines (Fig. 5E, F). When the assay was performed with PVR-co-expressing targets, we again only found a visible increase in degranulation when PVR was combined with B7H6 (Fig. 5G). Also, the enhancement of serially degranulating cells was much weaker in comparison to 2B4-co-engagement (Fig. 5H, I). Plotting all data for overall degranulation and fractions of serially degranulating cells revealed a positive statistical correlation with a coefficient of determination R^2^ = 0.7872 (Fig. S10). Upon closer examination, however, NK cells clearly and specifically diverged from the determined correlation depending on what receptor was engaged. NK cells incubated with B7H6– or CD48-expressing targets showed an over-representation of serially degranulating NK cells (Fig. S10). In contrast, co-incubation with PVR-, MICA-, or CD20-expressing targets was associated with a smaller fraction of serially degranulating cells. To validate the beneficial role of 2B4 co-stimulation, we performed time lapse microscopy of NK cells co-incubated with parental or CD48-expressing HeLa cells (Fig. 5J). Data showed that the fraction of NK cells killing one or multiple targets was clearly increased against CD48-expressing targets.

Overall, these data indicate that while the fraction of NK cells degranulating in serial fashion does correlate with the amount of overall degranulation, serial degranulation and killing can be specifically enhanced by co-stimulation of 2B4.

## Discussion

NK cell activity is regulated by the interplay of a plethora of inhibitory and activating receptors. Gaining insights into clear cause-and-effect relationships rising from specific ligand-receptor interactions is crucial for understanding NK cell activation and the advancement of NK cell-based therapies. There have been multiple studies performed with ligand-expressing *drosophila* S2 cells or plate-bound antibodies that focused on research questions similar to the ones investigated within this work (Bryceson et al., 2009, Bryceson et al., 2005, Bryceson et al., 2006b). These studies, however, focused largely on resting NK cells as effectors. In addition, optimal growth conditions for S2 cells such as culture medium and temperature differ significantly to those for human NK cells, making co-culture and particularly long-term assays under physiological conditions impossible, which severely limits the investigation of serial killing and cytokine release. We chose murine NIH3T3 cells as a cellular model system to enable co-incubation with human NK cells for long time spans without majorly disrupting cellular processes in targets. By employing pre-stimulated NK cells, properties of less potent co-activating ligand-receptor pairings could be studied more sensitively. With the development of therapeutic NK cell-based approaches in mind, this system more closely models the circumstances under which NK cells would be employed, e.g. in CAR-NK cell therapy. In this context, a major limitation so far is the exclusion of inhibitory signaling, which plays a pivotal role in tumor immune evasion (Seliger and Koehl, 2022). The systematic approach under which this work was conducted, however, easily allows for the inclusion of inhibitory ligands and numerous more combinations to gain even more insights into NK cell stimulation.

Engaging NKp30, NKG2D, or CD16 was sufficient to effectively trigger NK cell degranulation, cytokine secretion, and to induce target cell death. Discrepancies that were observed between ^51^Cr-release and impedance-based cytotoxic assays can be explained by methodical differences defining these two approaches. Since the latter relies on the adherent properties of target cells, it is likely that target cells overexpressing ICAM-1 on their surface caused faulty read-outs. Accordingly, ICAM-1-expressing target cells were shown to form a significantly larger number of prolonged conjugates with NK cells than any other employed ligand. Due to this property of LFA-1-ICAM-1-interactions, co-expression of ICAM-1 on target cells could be used as proxy to investigate the influence of adhesion on NK cell stimulation. While conjugate formation was enhanced as a result of ICAM-1-co-expression, degranulation, cytokine secretion and target cell death showed only minor increases compared to NK cells co-incubated with single ligand-expressing targets with the exception of CD20-expressing targets. Interestingly, combining ICAM-1 and MICA on target cells caused no increased NK cell degranulation but significantly increased cytotoxicity. LFA-1 has been described to support polarization of granules towards the immune synapse, implicating that granzyme release upon LFA-1-co-engagement is not increased globally but rather directed more efficiently against target cells (Barber et al., 2004). This hypothesis is further supported by the observed degranulation and target cell death induction of NK cells co-incubated with targets that were transfected exclusively with ICAM-1.

Our data confirm the downregulation of activating NK cell receptors upon exposure to their respective ligands that have been described elsewhere, including NKp30, NKG2D, and CD16. This downregulation was detectable on degranulating and non-degranulating NK cells, and the ligand-induced downregulation was only slightly reduced when activating signaling was blocked with dasatinib. Therefore, ligand specific downregulation of activating receptors seems to be upstream and independent of full NK cell activation and degranulation, creating the possibility of receptor-downregulation in an environment of high ligand expression without full NK cell activation and target cell killing.

CD16 and NKp46 were downregulated as a result of NK cell stimulation by different activating ligands. The downmodulation was fully dependent on NK cell activation as it could be blocked by dasatinib and it strongly correlated with the amount of degranulation. For CD16, receptor shedding has been described as a result of NK cell stimulation (Srpan et al., 2018). For NKp46 it is unclear how its downmodulation is regulated. So far, NKp46 has not been described to be shed upon NK cell activation and none of the employed target cell lines expressed ligands that verifiably bind to NKp46. A clear limitation of the employed assay in this regard is that different processes leading to receptor downregulation, e.g. internalization, shedding, or trogocytosis, cannot be identified or distinguished from one another. Investigating molecular mechanisms and functional consequences of this activation-dependent NKp46 downregulation will therefore require further research.

While clearly activation-dependent, downmodulations of CD16 and NKp46 were also observed on non-degranulation cells. This indicates that during the interaction between NK and target cells, activating signaling is induced but does not necessarily lead to degranulation. The reason for this is unclear. For CD16, a closer investigation of flow cytometry plots revealed that cells expressing particularly high levels (CD16^hi^) were not found in the population of non-degranulating cells (Fig. S11). This indicates that CD16^hi^ cells might be primed to degranulate, which would shift measured levels in non-degranulating cells artificially. While this was not observed to a similar extent for other receptors, it would be interesting to repeat this assay with NK cells sorted for high and low CD16 contents.

2B4 and DNAM-1 are described as co-stimulatory receptors, indicating that they only induce activity effectively when other receptors on the NK cell surface are engaged simultaneously (Sivori et al., 2000, Long et al., 2013). This is confirmed by our data showing lower degranulation and cytotoxicity against CD48-, PVR-, or Nectin-2-expressing target cells when compared to B7H6, MICA, or CD20. Co-expression of PVR on target cells caused only rare increases in NK cell activation. While degranulation and IFN-γ secretion were increased significantly when targets co-expressed B7H6, no effects were observed in combination with NKG2D– or CD16-stimulating ligands. It has been shown that NKG2D and DNAM-1 do not synergize with each other (Long et al., 2013), but specific relationships between DNAM-1 and CD16 or NCRs were largely overlooked so far. At first glance, our results seem to be in conflict with the classification of DNAM-1 as potent co-stimulatory receptor. However, PVR does not only bind to the co-stimulating receptor DNAM-1 but also to the inhibitory receptors TIGIT and CD96. This explains the downregulation of TIGIT, which we observed upon contact with PVR expressing targets. It further indicates that in a cellular environment, co-stimulatory properties of PVR are restrained due to competitive binding partners on the NK cell site.

Upon co-incubation with different targets, interactions with PVR but not Nectin-2 caused DNAM-1 and TIGIT levels on NK cells to decrease, revealing clear differences between these two ligands. While differences in the binding properties, including modes, affinities, and adverse homophilic binding between DNAM-1 and these two ligands have been described (Tahara-Hanaoka et al., 2004, Samanta and Almo, 2015, Wang et al., 2019), it is unclear why prolonged engagement of Nectin-2 did not induce downregulation of DNAM-1. Shedding light on this differential feedback regulation of DNAM-1 poses an interesting topic for further research.

When CD48 was co-expressed with any other ligand on target cells, multiple NK cell effector functions were enhanced significantly. Most obvious changes were observed in context of cytokine secretion and serial degranulation. While 2B4 has long been established as potent co-stimulatory receptor that can greatly boost NK cell activity in cooperation with various receptors (Urlaub et al., 2016, Bryceson et al., 2006b), its highly beneficial role in NK cell serial degranulation has not been described before. While we found a clear correlation between the fraction of NK cells degranulating in serial fashion and the amount of overall NK cell degranulation, our data also show that co-stimulation of 2B4 significantly increases the subpopulation of NK cells that degranulate against multiple targets. These results were confirmed by time lapse microscopy of NK cells with CD48-expressing HeLa cells by which the number of actual killing events per NK cell could be determined. Data clearly indicated that fractions of NK cells killing one or more targets was increased against CD48-expressing HeLa cells compared to parental controls. Considering the growing interest and research in NK cell-based cancer immunotherapies (Dos Reis et al., 2025, Myers and Miller, 2021), these data highlight the potential benefits achievable by including 2B4-based signaling in therapeutical approaches. Significantly, a recent study on GPC3-directed CAR-NK cells could show enhanced cytotoxicity against target cancer cells upon inclusion of the intracellular domain of 2B4 (Huang et al., 2020).

Taken together, this study provides a robust foundation highlighting the specific impact of distinct human NK cell ligand-receptor interactions on NK cell effector functions. While several aspects will need to be investigated in more detail in future projects, our results may already prove to be vital in the development of NK cell-focused therapeutic approaches.

## Author contributions

LK and CW conceptualized the study. LK designed and performed the experiments. MSar performed the conjugate formation assays shown in Fig. 1A/B, 2A/B. JN performed and analyzed the microscopy-based serial killing assay shown in Fig. 5J. MSan generated the expression vectors employed for transfections. SW and MC sorted transfected cells for purity. LK managed, analyzed, and visualized all data. CW administrated and supervised the study. LK and CW wrote the manuscript. The results and the original draft were reviewed and discussed by all authors.

## Acknowledgements

We would like to thank all members of the Watzl Immunology Lab for the fruitful discussions concerning this project and all kinds of banter held over lunch that motivated the authors to keep going. We further thank Annika Winterberg for her contributions to this project. This work was supported by internal funds of the Leibniz Research Centre for Working Environment and Human Factors (IfADo).

