## Supplemental figures for "2B4 co-engagement promotes serial degranulation and killing by human NK cells"

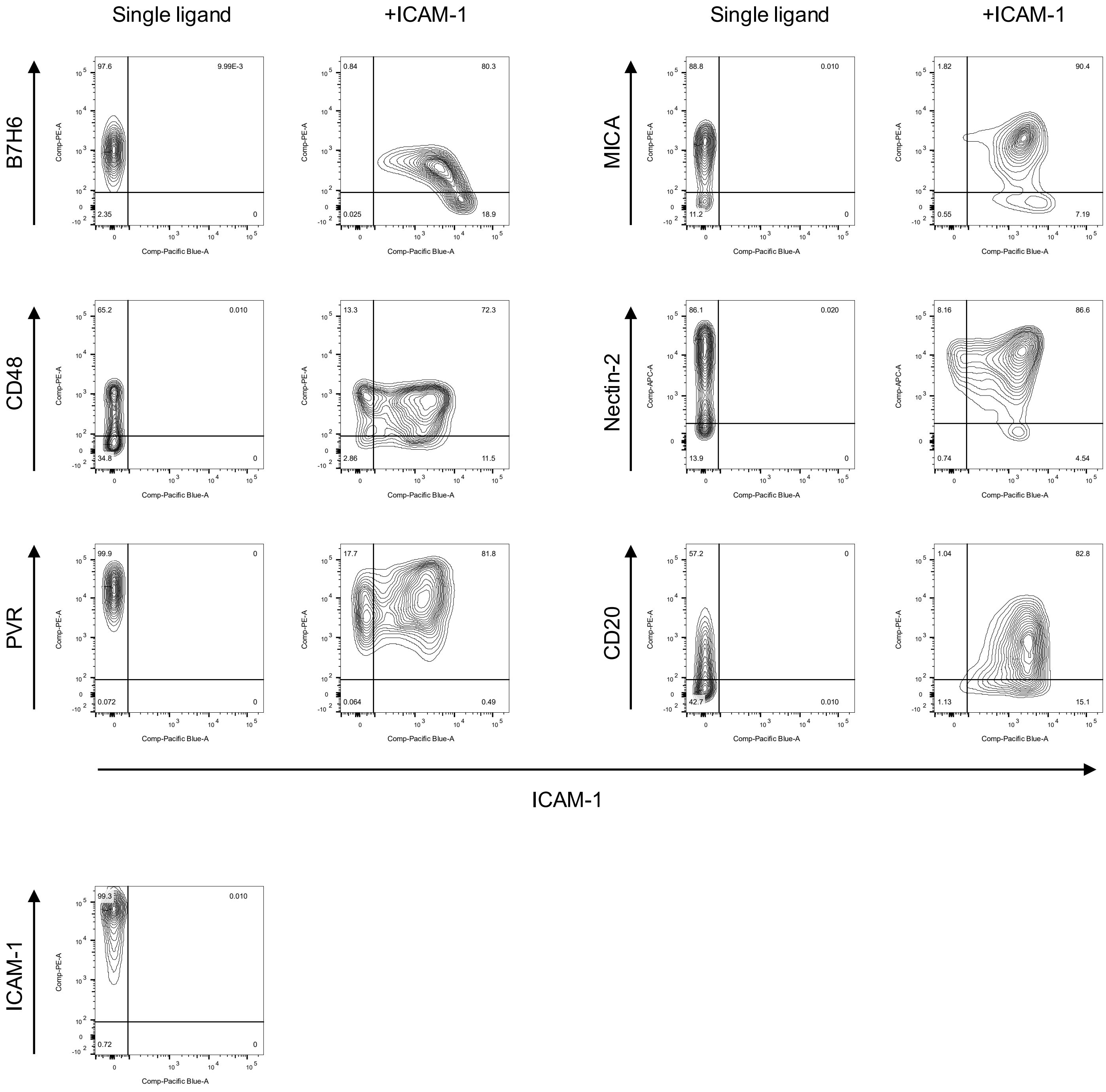


**Figure S1:** Expression of human NK cell ligands in respective transfected NIH3T3 cells depicted in flow cytometry density plots. Left column: cells transfected with a single ligand. Right column: cells transfected with ICAM-1 in addition to another respective ligand. X-axes show ICAM-1 levels; Y-axes show levels of respective single ligands.

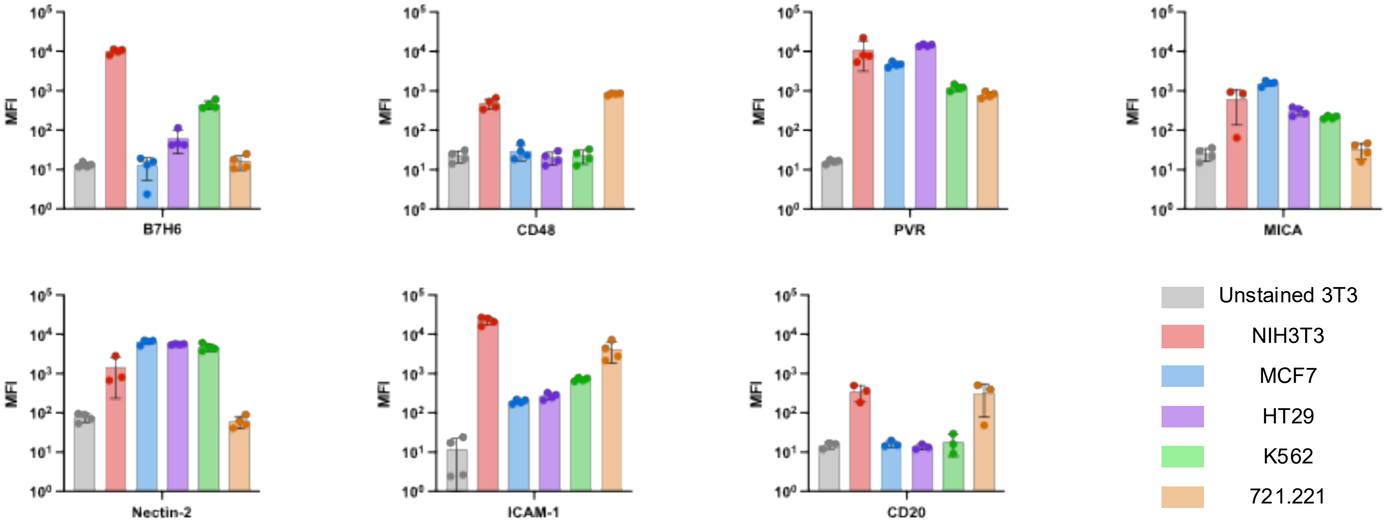


**Figure S2:** Comparison of relative expression levels determined by flow cytometry of human NK cell ligands in transfected NIH3T3 cells and a selection of commonly employed human cancer cell lines; n = 5.

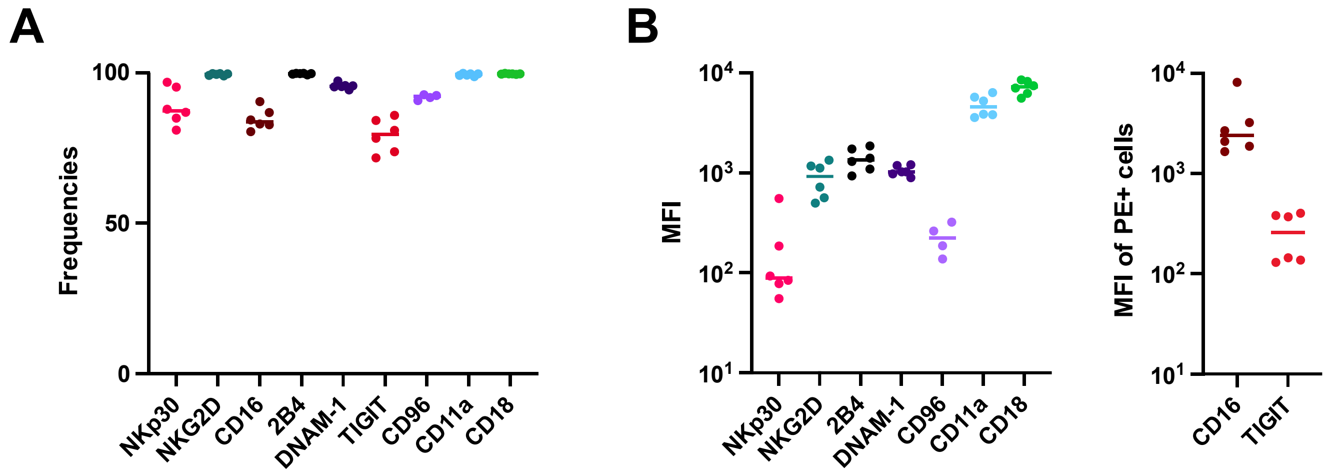


**Figure S3:** Variability of relative expression levels of NK cell receptors between NK cell donors determined by flow cytometry; n = 4-6. Figures show quantifications of **A** frequencies of receptor positive cells and **B** MFIs of respective receptors in all cells (left) or in receptor positive cells (right).

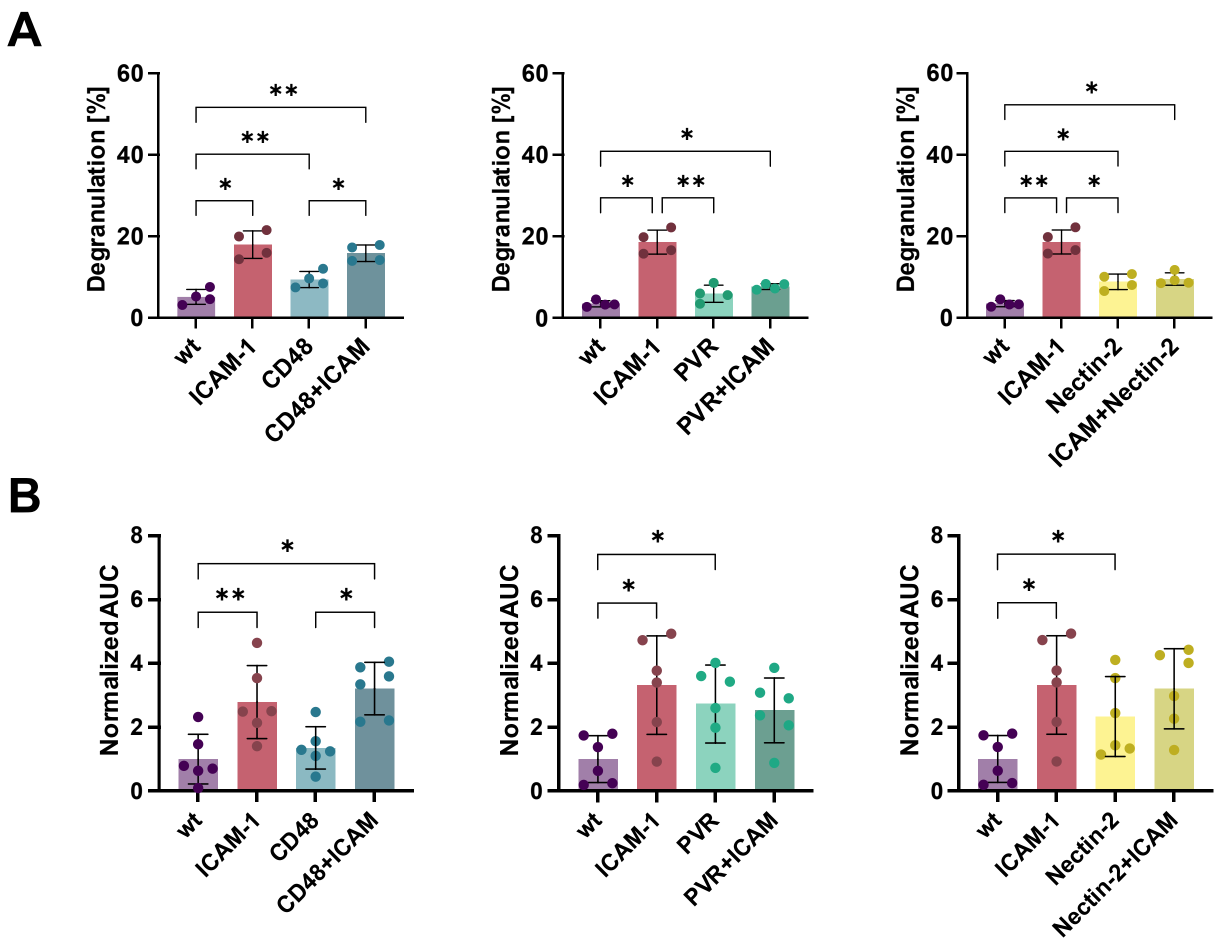


**Figure S4:** Results of **A** flow cytometry-based degranulation (n = 4) and **B** $^{51}$Cr-release cytotoxicity (n = 6) assays against target cells co-expressing ICAM-1 and CD48, PVR, or Nectin-2, respectively. *Data information*: One-Way ANOVA with Šidák correction. *$P \le 0.05$, **$P \le 0.01$. Data presented as mean ± SD.

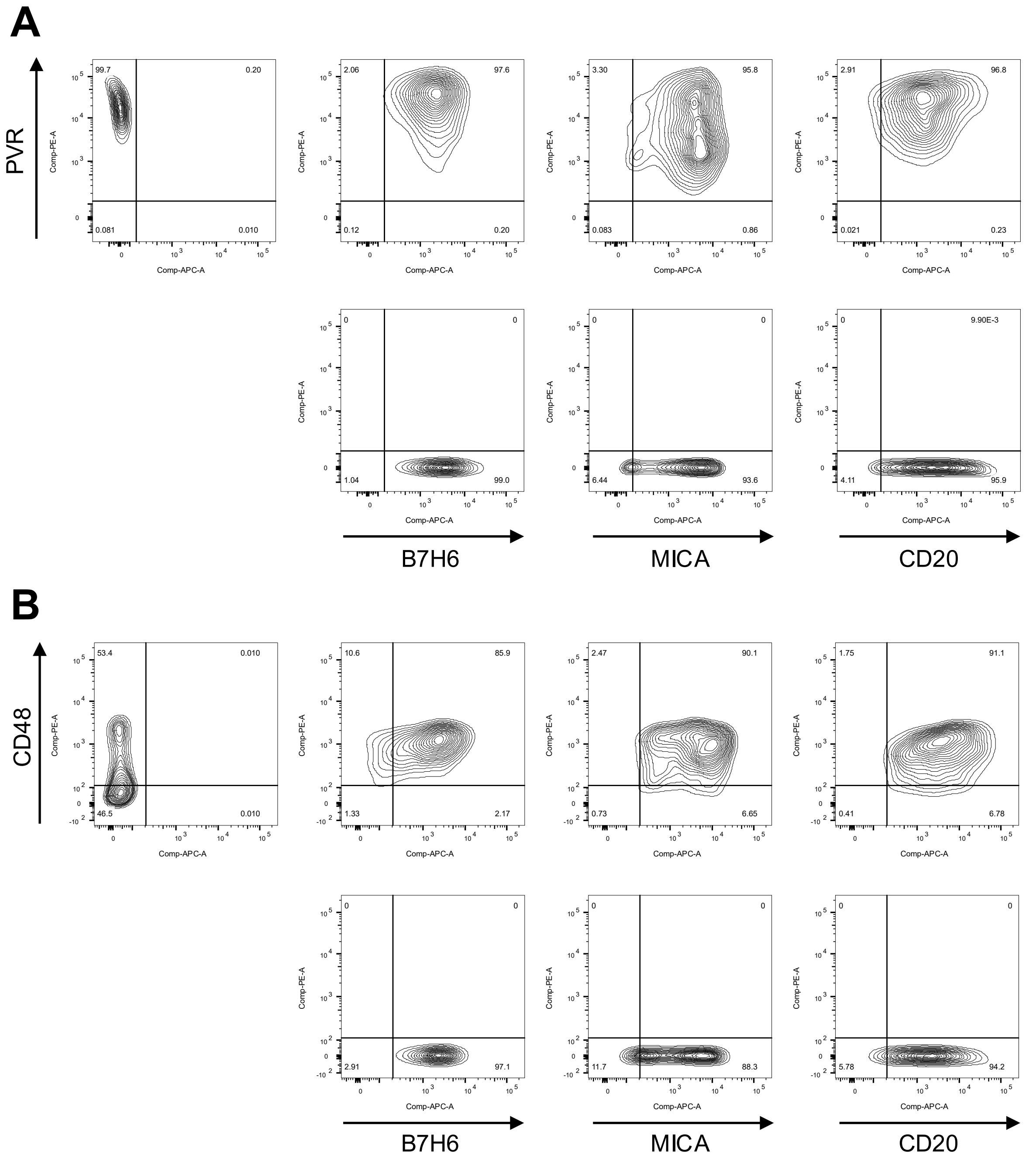


**Figure S5:** Comparison of expression of human NK cell ligands in NIH3T3 cells transfected with one or a combination of two ligands depicted in flow cytometry density plots. X-axes show levels of respective single ligands; Y-axes show levels of **A** PVR or **B** CD48.

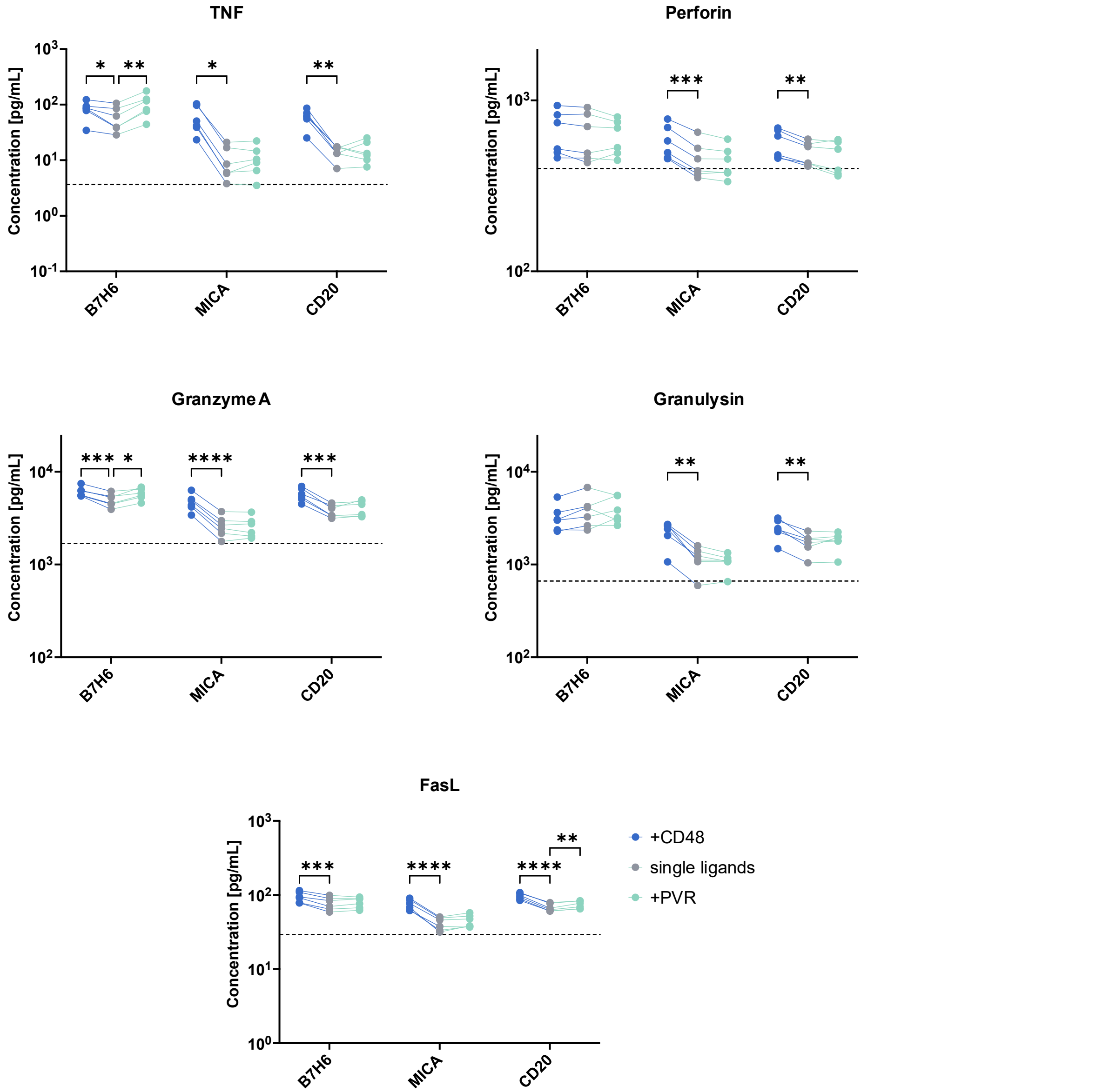


**Figure S6:** Secretion levels of TNF, Perforin, GrzA, Granulysin, and FasL after overnight co-incubation of NK cells with CD48- or PVR-co-expressing targets at an E/T ratio of 1:1. Dashed lines indicate means of NK cell controls without targets; n = 6. *Data information*: Two-Way ANOVA with Dunnet correction. $*P \leq 0.05$, $**P \leq 0.01$, $***P \leq 0.001$, $****P \leq 0.0001$.

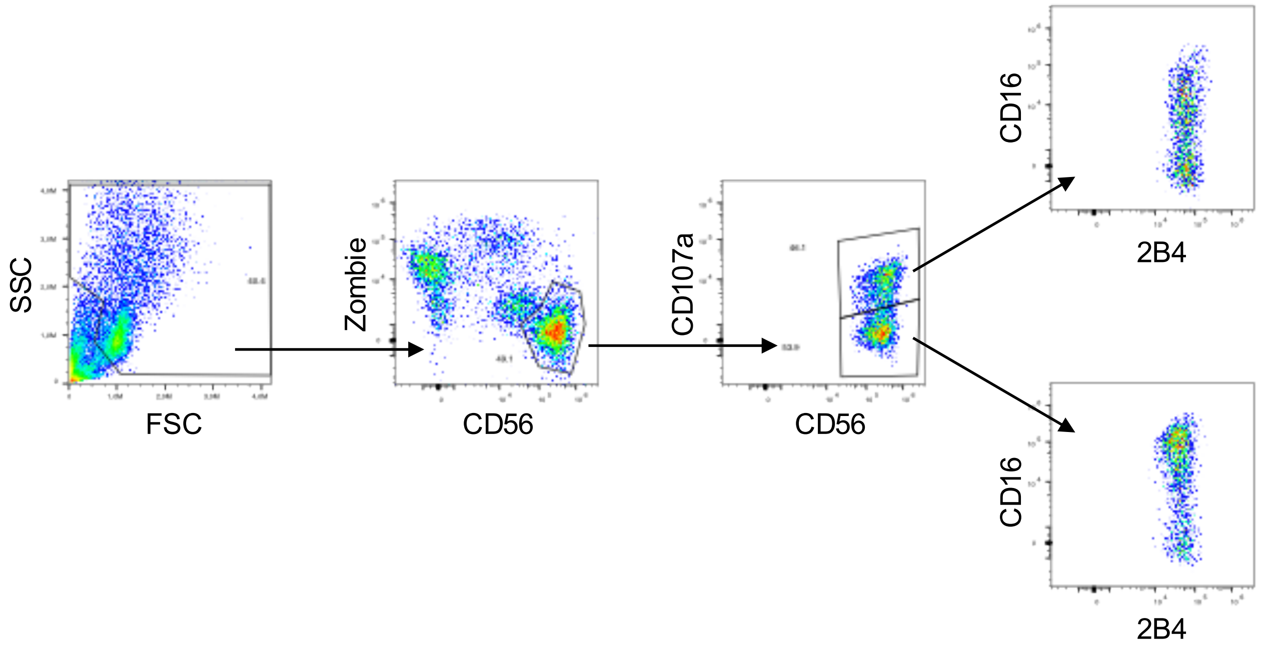


**Figure S7:** Gating strategy employed for the determination of degranulation-dependent changes in NK cell receptor levels after co-incubation with different target cell lines.

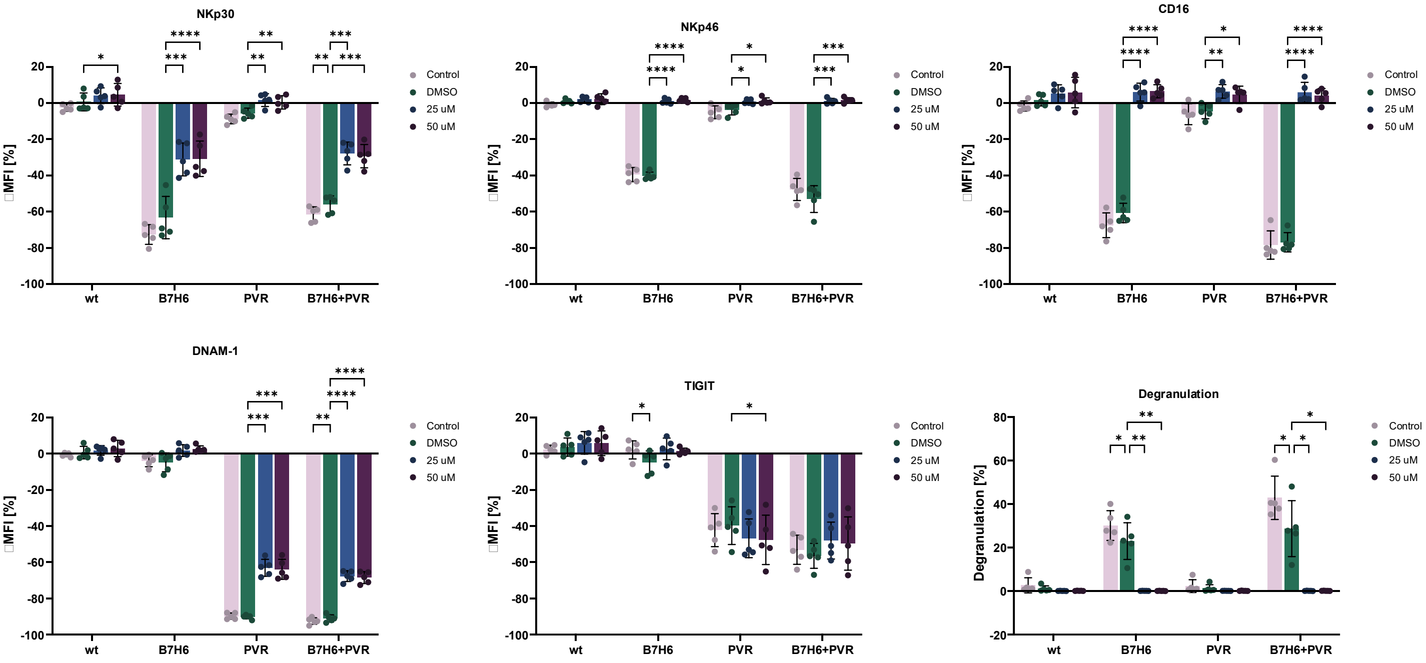


**Figure S8:** Changes in expression levels of different NK cell receptors after treatment with tyrosine kinase inhibitor dasatinib and subsequent co-incubation with different target cells; n = 5. *Data information :* Two-Way ANOVA with Dunnet correction. *$P \leq 0.05$, **$P \leq 0.01$, ***$P \leq 0.001$, ****$P \leq 0.0001$. Data presented as mean ± SD.

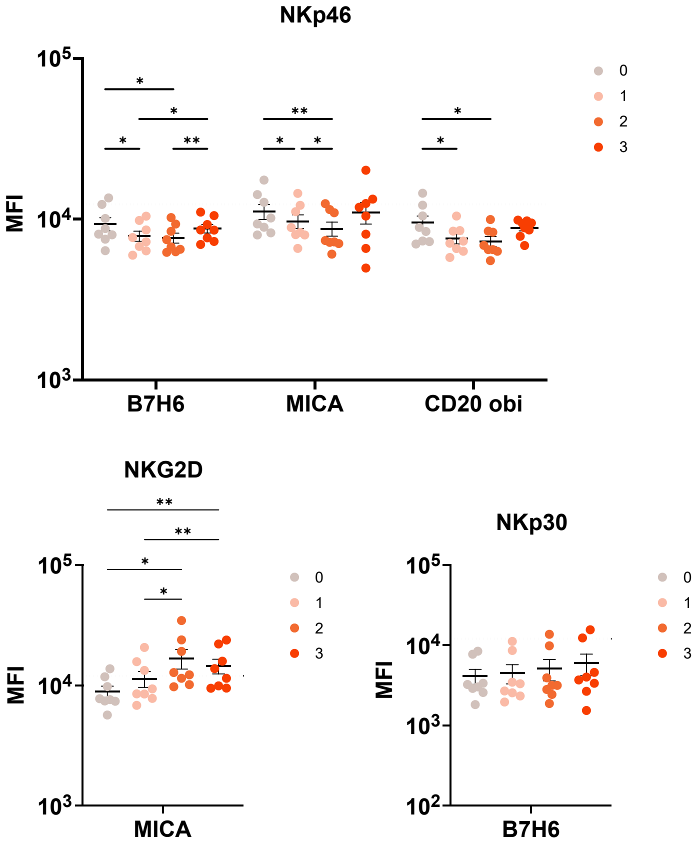


**Figure S9:** NK cell receptor expression levels after co-incubation with different targets depending on the number of degranulation events per NK cell. Dashed lines indicate expression levels of NK controls without targets; n = 8. *Data information:* One-Way (NKp30, NKG2D) or Two-Way (NKp46) ANOVA with Tukey correction. *$P \leq 0.05$, **$P \leq 0.01$. Data presented as mean ± SEM.

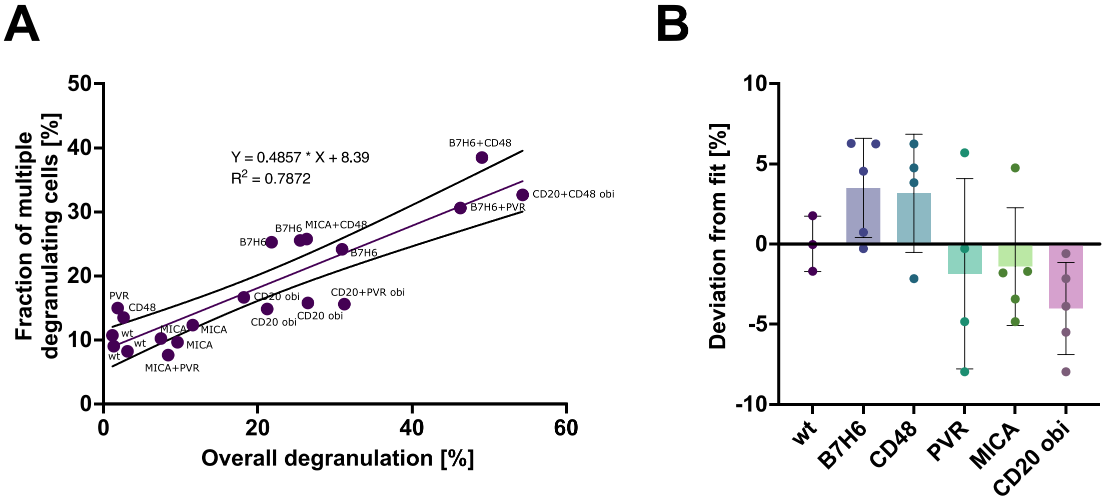


**Figure S10: A** Correlation of overall degranulation against fractions of serially degranulating cells. Data points indicate means of NK cells co-incubated with targets expressing respective ligands. There are three data points each for wt, B7H6, MICA, and CD20 obi because assays were conducted in three separate setups: with targets expressing single ligands, targets co-expressing CD48, and targets co-expressing PVR. **B** Deviations of the fractions of serially degranulating cells from the linear fit introduced in panel A. *Data information:* Linear regression with 95 % confidence bands (A). Data presented as mean ± SD (B).

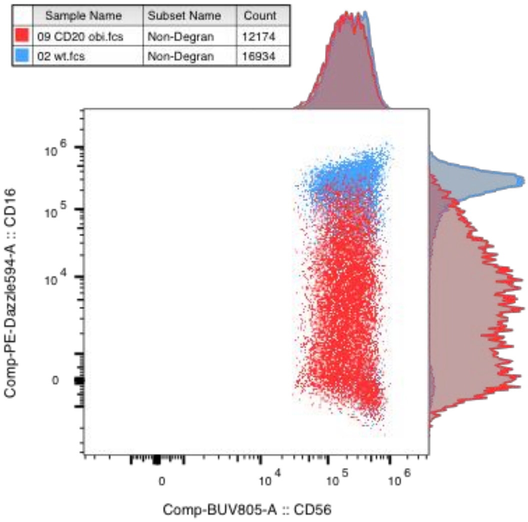


**Figure S11:** Representative flow cytometry plot showing the expression of CD16 in NK cells that did not degranulate after co-incubation with wild type (blue) or CD20-expressing target cells (red) treated with obinutuzumab.
